# Light Harvesting Complex II Resists Non-Bilayer Lipid-Induced Polymorphism in Plant Thylakoid Membranes via Lipid Redistribution

**DOI:** 10.1101/2024.11.15.623718

**Authors:** Avinash Garg, Ananya Debnath

## Abstract

The plant thylakoid membrane hosting the light-harvesting complex (LHCII) is the site of oxygenic photosynthesis. Contrary to the earlier consensus of a protein-driven single lamellar phase of the thylakoid despite containing 40% non-bilayer-forming lipids, recent experiments confirm the polymorphic state of the functional thylakoid. What, then, is the origin of this polymorphism, and what factors control it? The current Letter addresses the question using a total of 617.8 *μ*s long coarse-grained simulations of thylakoids with and without LHCII and varying concentrations of non-bilayer lipids using Martini-2.2 and 3.0 at 323 K. The LHCII redistributes the non-bilayer lipids into its annular region, increases the bending modulus and the stalk formation free energy, reduces the non-zero mean curvature propensity, and resists the polymorphism these lipids promote. The thermodynamic trade-off between non-bilayer lipids and LHCII dictates the degree of nanoscopic curvature leading to the polymorphism crucial for non-photochemical quenching under excess heat conditions.

**Graphical TOC Entry:** 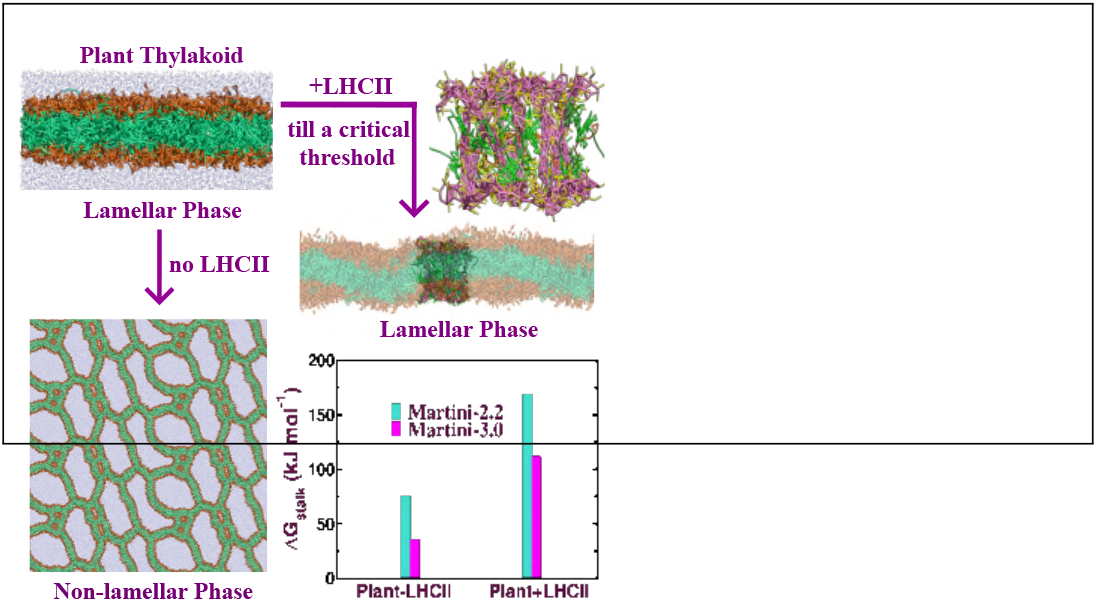

---

Plant chloroplasts comprise thylakoid membranes, a unique internal superstructure having stacked grana connected with stromal lamellae. The thylakoids provide a matrix for the pigmented LHCII protein trimer stabilized by the trimerization motif^1^ actively involved in the light-dependent photosynthetic processes. ^2^ On excess illumination, the LHCII regulates energy transport by non-photochemical quenching (NPQ) by conformational changes. ^3^ The dynamical structure or the phase of the thylakoid membrane plays a pivotal role in regulating photosynthesis in diverse environmental conditions^4^ by possible rearrangements^5^ connected to polymorphism of lamellar phase, isotropic phase or inverted hexagonal phase. The dynamic grana stacking of the thylakoid is mostly determined by phosphorylation and adapts to different light conditions.^6^ The dynamical aggregation of chlorophyll induces lamellar to non-lamellar phase transition in thylakoids. ^7^ The specific conformation of the LHCII and the compositions of the thylakoid membranes are indispensable for stabilizing the photosynthetic machinery^8^ and enabling the NPQ.^9^ However, the mechanisms by which LHCII modulates thylakoid topology and its relationship to lightharvesting and NPQ remain pressing questions that need to be addressed.

Unlike other biological membranes, the thylakoid membrane primarily consists of conserved glycoglycerolipids named monogalactosyldiacylglycerol (MGDG, MGDT), digalactosyldiacylglycerol (DGDG, DGDT), sulfoquinovosyldiacylglycerol (SQDG) and phospholipids of phosphatidylglycerol (PG, PT). 40% of the lipids in thylakoid membranes are MGDG and MGDT, non-bilayer forming lipids that can undergo lipid polymorphism and form various non-lamellar phases such as isotropic, inverted hexagonal (H_*II*_), and cubic phases.^10^ In contrast, DGDG with a cylindrical shape can assemble into a lamellar phase. SQDG and PG have anionic head groups with acidic properties affecting the interactions and stability of the membrane and the protein.^11^ Under physiologically relevant conditions, the thylakoid lipid mixture cannot produce lamellar phases without proteins but adopts various non-bilayer forms.^12^ This is consistent with the results from coarse-grained (CG) molecular dynamics simulations of the thylakoid membrane without the LHCII at a critical hydration level.^8^ Remarkably, MGDG self-assembles into a large lamellar aggregate in the presence of LHCII.^13^ MGDG helps in the structural association of LHCII through lateral pressure^14^ and prevents LHCII from unfolding due to the steric matching. ^15^ Earlier there was a consensus that the thylakoid membrane remains in a single lamellar phase driven by the proteins while the stack formation is facilitated by the non-bilayer-forming lipids.^16^ Although the thylakoid consists of ∼ 40% non-bilayer lipids, the origin of the single lamellar phase of the thylakoid remains a paradox. On the contrary, recent experiments suggest that the fully functional thylakoid membrane contains a H_*II*_ phase, two isotropic phases^17^ and a lamellar phase.^18^ The thylakoids host key enzymes of the xanthophyll cycle, violaxanthin de-epoxidase, VDE^19^ to help excess energy dissipation in NPQ. ^20^ The non-bilayer forming lipids, MGDG and MGDT, which are located near the LHCII,^15^ significantly enhance the VDE activation in the H_*II*_ phase of the thylakoid.^20^ VDE activity is influenced by temperature and pH-dependent reorganizations of non-lamellar phases, ^21^ thus controlling NPQ. This suggests that activation of the xanthophyll cycle for NPQ can be triggered or modulated by thylakoid membrane polymorphism. Despite evidence linking lipid polymorphism to light-harvesting and NPQ, no systematic analysis is performed to quantify the key determinants of functional thylakoid polymorphism which requires urgent attention. Thus, the present letter aims to elucidate the molecular origin of functional thylakoid polymorphism, a topic that has been contentious in previous studies. A total of 617.8 *μ*s unbiased and biased CG MD simulations are carried out for the membranes with 0 − 59% of non-bilayer lipids along with the remaining lipids of the plant thylakoid and LHCII with chlorophyll and lutein embedded in thylakoid at 323 K using Martini-2.2^22^ and Martini-3.0.^23^ In the absence of neoxanthin and violaxanthin, their binding sites are mostly occupied with lipids, DGDT, MGDT, and SQDG. The schematic representation of the chloroplast, the CG structure of the bilayer, and the mapping and composition of the plant thylakoid membrane are presented in Figure 1. The simulation details and computational methods are discussed in the Supporting Information (SI) where Table S1 of the SI mentions the system details of unbiased runs.

**Figure 1:**
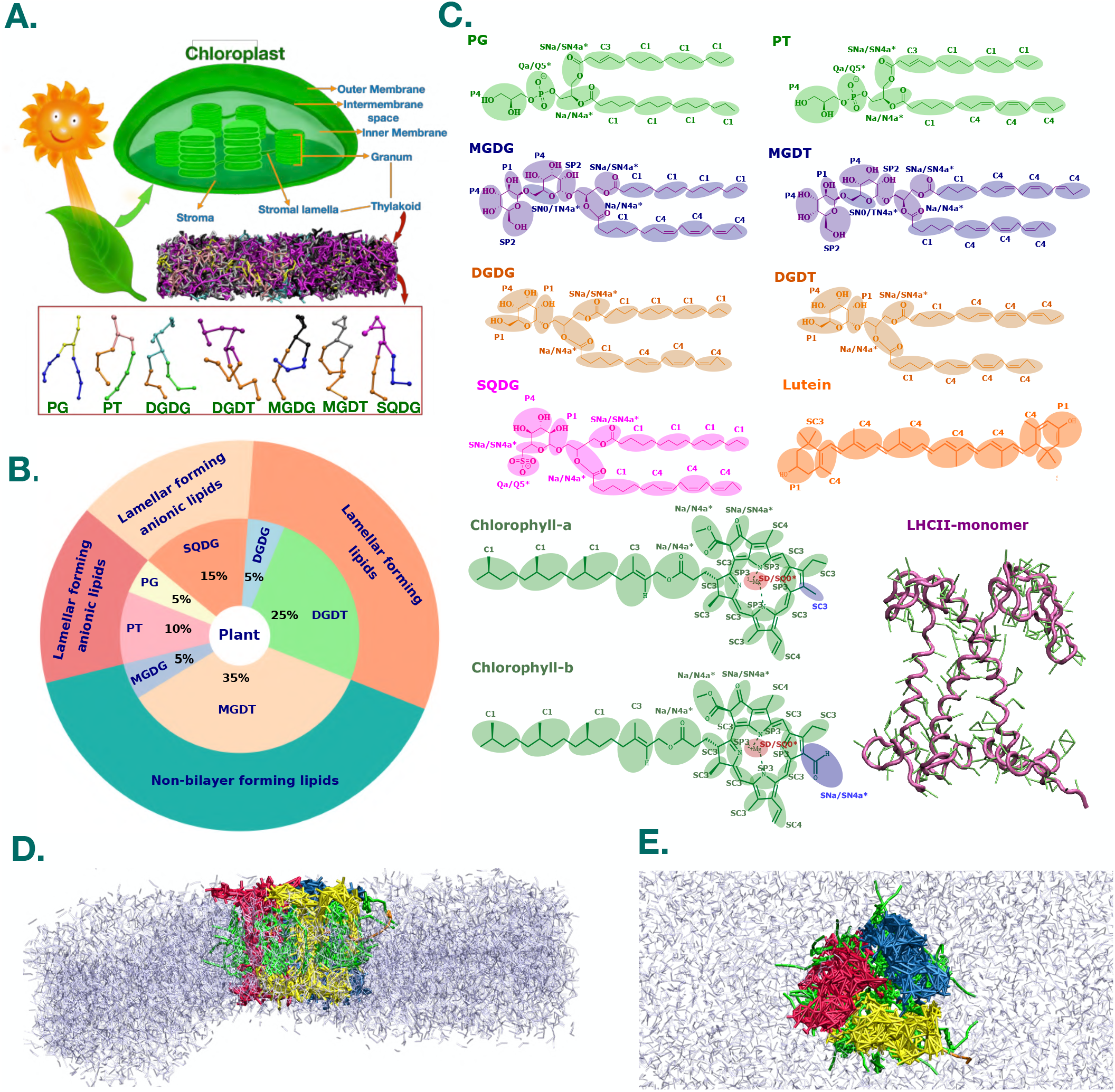
(A) Schematic representation of the Chloroplast and CG respresentations of seven lipids comprising the thylakoid membrane. (B) Pie chart showing the composition of the plant thylakoid. (C) CG beads superimposed on the AA chemical formula of seven lipids (PG, PT, MGDG, MGDT, DGDG, DGDT, SQDG) and the cofactors of LHCII: lutein, chlorophyll-a and b. CG bead types of Martini-2.2 and Martini-3.0 are mentioned near the CG beads shown in transparent spheres. The starred (*) CG beads represent the change in bead type from Martini-2.2 to Martini-3.0. The CG structure of the LHCII monomer in tube representation: backbone beads in magenta and side chains in green. (D) Side and (E) top view of the LHCII trimer embedded in the thylakoid membrane. Three monomers of the trimer in blue, red, and yellow; lipids in ice-blue; lutein in orange; chlorophyll-a and b in green.

A lamellar to non-lamellar phase transformation is identified from a drastic reduction in area per lipid, *a*_*h*_, within 5 *μ*s simulations of stacked bilayers without the LHCII having non-bilayer lipids from 35 − 59% including the plant composition using anisotropic pressure coupling (Figure S1 of the SI). No changes in *a*_*h*_ are observed for the thylakoid membranes without the non-bilayer lipids and the plant thylakoids with the LHCII even for 10 *μ*s NPT run-length suggesting no polymorphism. Single or stacked membranes without LHCII using semi-isotropic pressure coupling do not exhibit any phase transition in MARTINI-2.2. However, the membranes without the LHCII undergo phase transitions using semi-isotropic pressure coupling in MARTINI-3.0. The thickness of the plant thylakoid membrane from our simulations (Figure S2 of the SI) is similar to that of the thylakoid with and without PsbS upon illumination.^24^ The unbiased simulations of the bilayers with varying concentrations of non-bilayer lipids using Martini-2.2 and 3.0 and anisotropic pressure coupling demonstrate a lamellar to non-lamellar phase transformation at different time scales as seen in Figure 2. The fused regions are occupied mostly by non-bilayer lipids and these regions slowly convert to the non-lamellar cubic or H_*II*_ phase within the initial 5 *μ*s. The potential impact of GROMACS parameters on our results has been discussed in the context of^25^ in the SI (Figure S3-S5).

**Figure 2:**
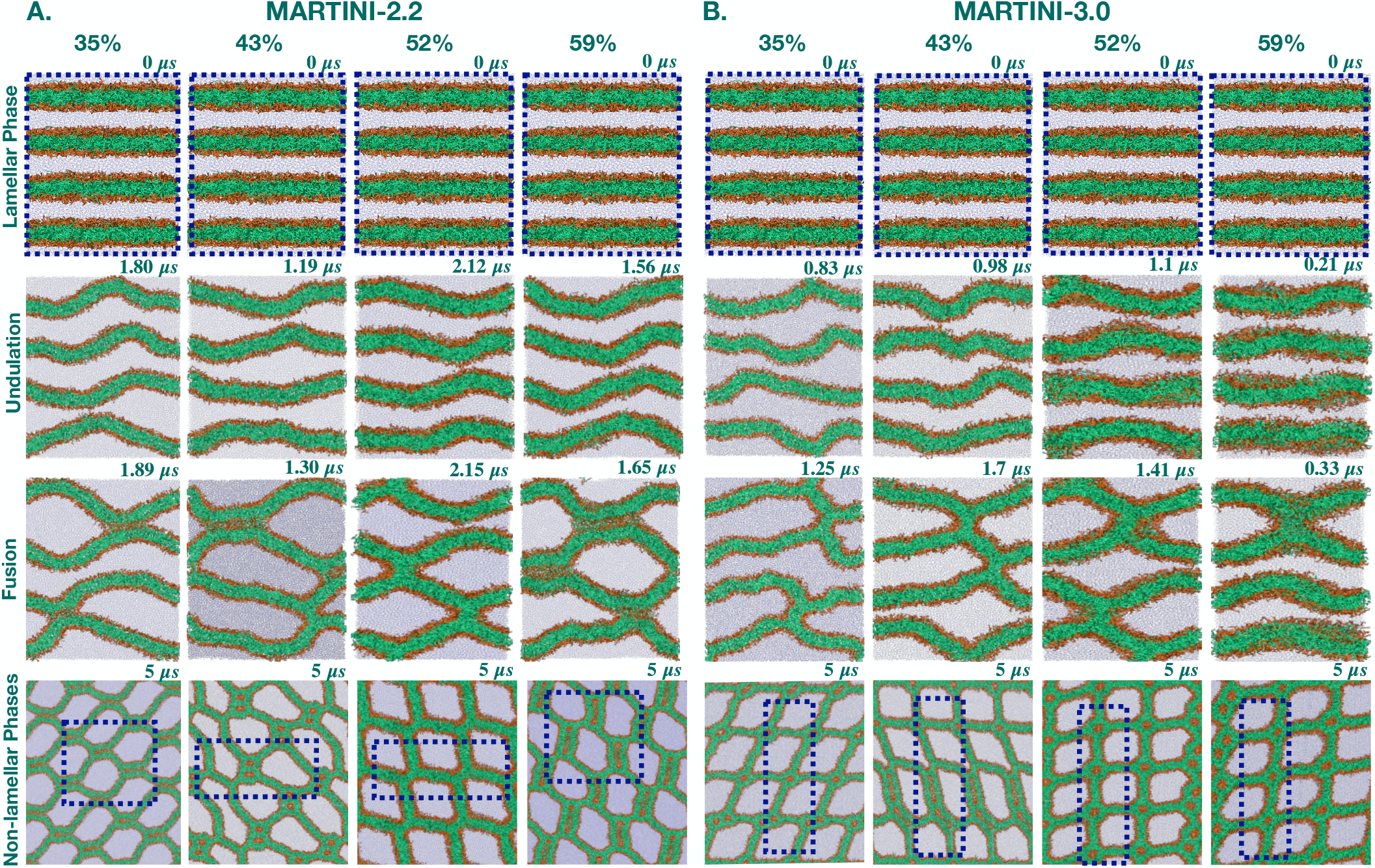
The lamellar to non-lamellar phase transformation (inverted hexagonal for 35% and cubic for the remaining concentrations in MARTINI-2.2 and cubic for all concentrations in MARTINI3.0) of the bilayers via undulation and fusion using unbiased molecular dynamics simulations with anisotropic pressure coupling for (A) Martini-2.2 and (B) the Martini-3.0. The bilayers have 35%, 43%, 52%, and 59% non-bilayer lipids along with the remaining thylakoid compositions. During simulations, the bilayers quickly undergo undulations bringing the nearest membrane surfaces close. As a result, the intermediate water molecules go away, and the bilayer surfaces fuse within the initial 1.3 − 2.2 *μ*s for the Martini-2.2 and within the initial 0.33 − 1.7 *μ*s for the Martini-3.0. A dotted blue box represents the unit cell. Color code: lipid head groups: orange; tails: green; water: ice-blue points.

No phase transformation is observed for the bilayer without the non-bilayer lipids (see Figure 3A) using Martini-2.2 and 3.0 within initial 5 *μ*s. A lamellar to non-lamellar H_*II*_ or cubic phase transformation is again observed for the plant thylakoid membrane within initial 5 *μ*s for both Martini-2.2 and 3.0 via undulation and fusion similar to the other bilayers with non-bilayer lipids (Figure 3B). This aligns with previous observations^8^ of the CG plant membrane undergoing a spontaneous transition from a lamellar to an inverted hexagonal phase at a critical hydration level of 9 CG water beads per lipid at 293 K using Martini-2.2 with semi-isotropic pressure coupling unbiased MD simulations. Remarkably, no fusion is observed for the plant thylakoid bilayer in the presence of the LHCII as shown in Figure 3C. Thus, the lamellar to non-lamellar phase transformation or lipid polymorphism is triggered by the nonbilayer lipids but resisted by the LHCII. Earlier experiments show that LHCII forms lamellar structures when mixed with the non-bilayer lipids, MGDG, at ratios similar to those found in thylakoids.^13^ Our results demonstrate that the LHCII withstands non-lamellar phases and promotes bilayer organization, even when nonbilayer lipids are present.

**Figure 3:**
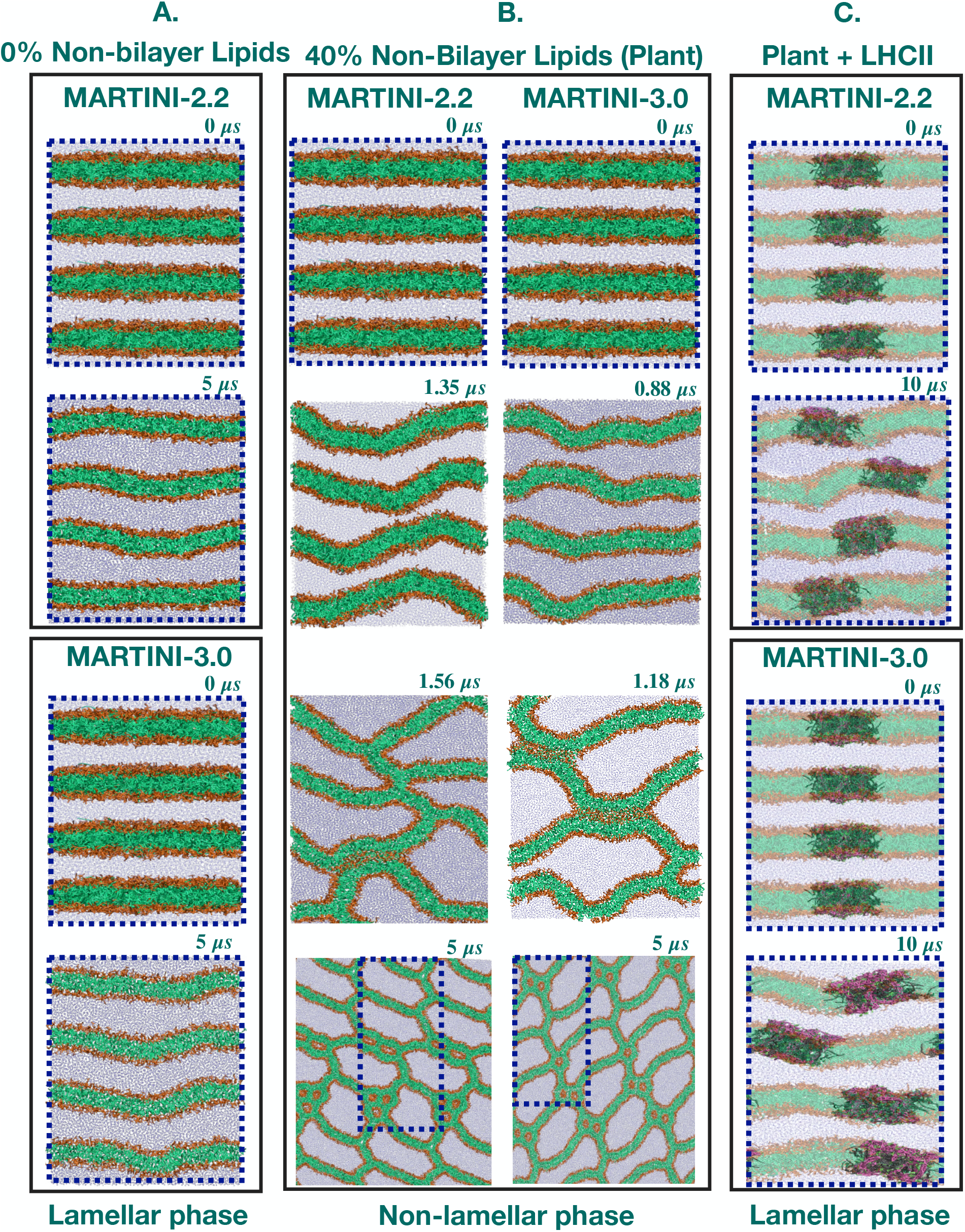
Snapshots from unbiased simulations using anisotropic pressure coupling with Martini-2.2 and 3.0 of (A) the thylakoid membrane without non-bilayer lipids showing no phase transformation. (B) Without LHCII, the plant thylakoid membrane undergoes a lamellar to non-lamellar phase transformation (inverted hexagonal in MARTINI-2.2 or cubic in MARTINI-3.0) via undulation and fusion. (C) The LHCII in plant thylakoid membrane resists non-lamellar phase transformation. A dotted blue box represents the unit cell. Color code: lipid headgroups: orange; tails: green; water: ice-blue points; LHCII trimer: mauve. The lipids in the presence of the LHCII are presented in transparent for clarity.

The phase transformation of the plant thylakoid membrane is quantified from membrane mean curvature, *H*, as follows,

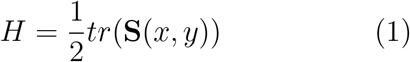

calculated over 10 − 20 replicas of systems (see methodology and Table S2 of the SI). The higher probability of non-zero mean curvature in plant thylakoids without the LHCII suggests a strong likelihood of transitioning from lamellar to non-lamellar phases lacking LHCII (Figure 4A). Nevertheless, when LHCII is present, the probabilities of the non-zero mean curvature reduce while the probabilities of the zero mean curvature increase (Figure 4B). This suggests that LHCII increases the likelihood of maintaining the thylakoid’s lamellar phase, resisting its transformation into a non-lamellar phase.

**Figure 4:**
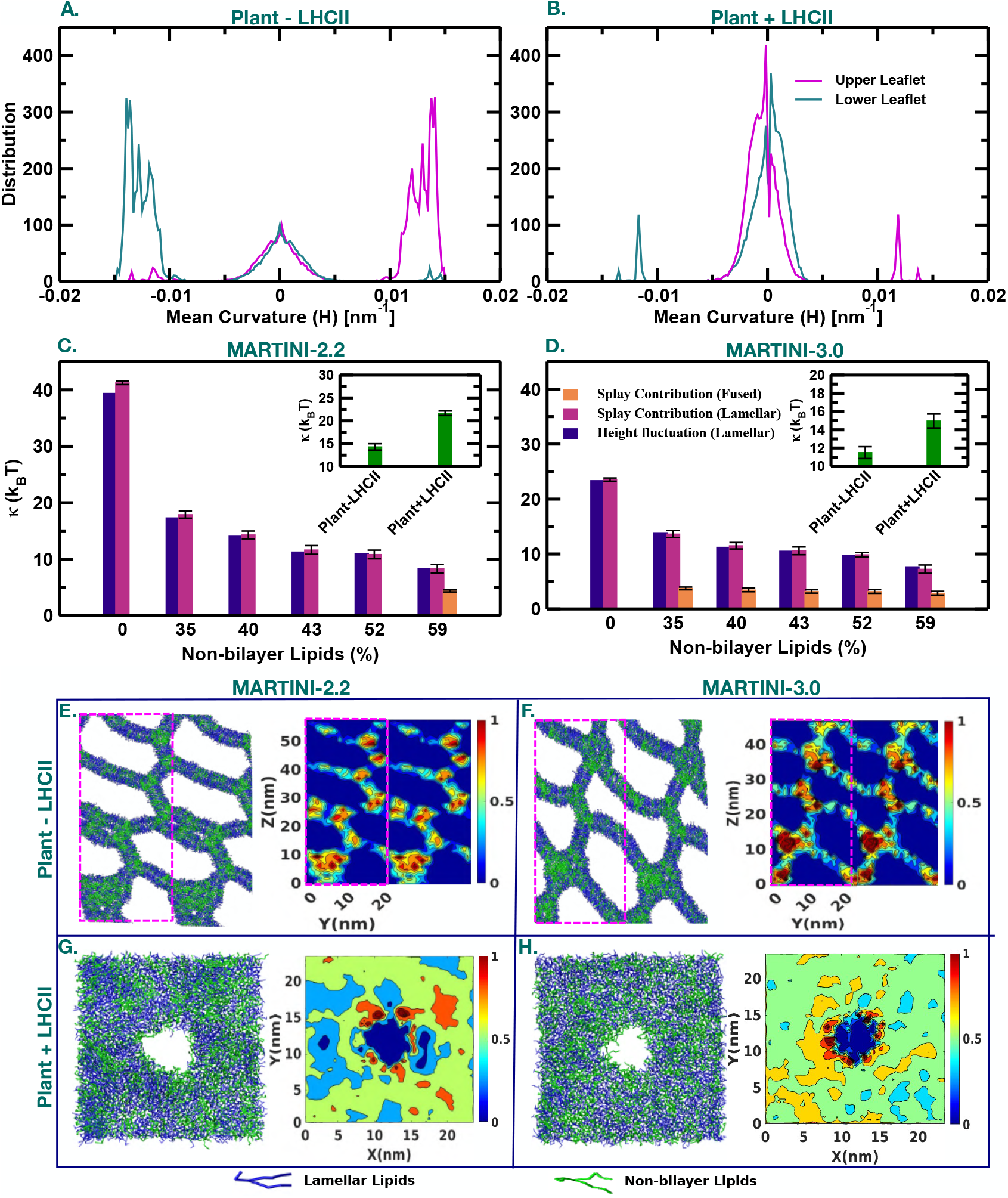
Distributions of the mean curvature, *H*, of the plant thylakoid membrane (A) without and (B) with the LHCII using MARTINI-3.0. The higher probability of the positive and negative *H* of the upper and lower leaflets, respectively compared to the zero mean curvature in (A) demonstrate the more probable non-lamellar phase over the lamellar phase in the absence of the LHCII. The higher probability of zero *H* over the non-zero ones in (B) demonstrates that the lamellar phase is more probable over the non-lamellar phase in the presence of the LHCII. Bending modulus, *κ*, of the bilayers with varying concentrations of non-bilayer lipids (0 − 59%) using (C) Martini-2.2 and (D) Martini-3.0 from the height fluctuation method for the lamellar phase (blue bars), and the splay contributions for the lamellar phase (magenta bars) and the fused phase (orange bars). A comparison of the *κ* of the plant thylakoid in the absence and presence of the LHCII is shown in the inset. Snapshots of the plant thylakoid bilayer with non-bilayer lipid density maps averaged over 5000 frames in (E) and (F) without LHCII using anisotropic pressure coupling and; (G) and (H) with LHCII using Martini-2.2 and Martini-3.0. Color code: lamellar lipids: blue; non-bilayer lipids: green; pink dotted box: unit cell. The density of non-bilayer lipids is the highest in the fused region and the annular region of the LHCII in the absence and presence of LHCII respectively. Water and LHCII have not been shown for clarity. All data except in E and F are obtained using semi-isotropic pressure coupling.

The resistance of the bilayers for the phase transformation is estimated through the bending modulus, *κ*^26^ calculated for the tensionless bilayers from the intensity spectra,

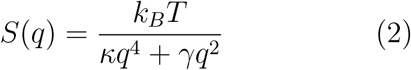

at the small *γ*, interfacial tension and small *q*, wave vector limit^27^ from height fluctuations of bilayer leaflets^28^ (see Figure S6 and Table S3 of the SI). Alternatively, *κ* of the multicomponent bilayer is calculated from the monolayer bending modulus *k*_*m*_ = *κ/*2 by considering the molecule pairwise splay contributions^29^ using

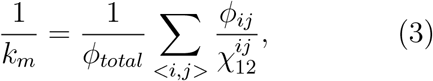

where 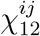 is the splay modulus (see the methodology and Figure S7-S9 in the SI) for the *ij*^*th*^ pair type and *ϕ*_*ij*_ is the number of near-neighboring *ij* encounter pairs, *ϕ*_*total*_ is the total number of encounters for all pairwise contributions for which the splay is calculated. Our methods are validated using the preequilibrated^30^ AA DMPC bilayer as mentioned in Table S4 of the SI. The *κ* of the thylakoid membranes decreases with increased concentration of non-bilayer lipids for both Martini-2.2 and 3.0 using both approaches (Figure 4 C and D, Table S5 of the SI). Thus the non-bilayer lipid reduces rigidity against bending or deformation which facilitates the lamellar to nonlamellar phase transformation. The reduction in *κ* with increasing non-bilayer lipids obtained for the fused state is marginal compared to that for the lamellar state, possibly due to the similar splay contributions in all fused states. The LHCII resists the lamellar to non-lamellar phase transformation by increasing the *κ* for Martini-2.2 and 3.0 (inset of Figure 4 C and D) which reduces the membrane fusion probability.

To estimate the stalk formation free energy, a chain reaction coordinate, *ξ*_*ch*_, is introduced using a membrane-connecting cylinder decomposed into *N*_*s*_ slices (Figure 5A-C) filled up with lipid beads, following the reference^31^ using umbrella sampling simulations (see Table S6 and section 1.6 of the SI for details). The histograms in Figures S10 and S11 of the SI confirm a good sampling of the *ξ*_*ch*_ and no hysteresis respectively. The stalk formation free energy, Δ*G*_*stalk*_, obtained from the difference in the PMF of the flat and stalk membrane is the highest (Table S7 of the SI) for the bilayer without non-bilayer lipids from Martini-2.2 and 3.0. For both Martini-2.2 and 3.0, the Δ*G*_*stalk*_ decreases with increasing concentration of the non-bilayer lipid concentrations (inset of Figure 5 D and E). The stalk formation becomes spontaneous for the bilayers with non-bilayer lipid concentrations of 52 − 59% for Martini-3.0 (Table S7 of the SI) since the Δ*G*_*stalk*_ is reduced by ∼ 20 − 40 kJ mol^*−*1^ from Martini-2.2 to 3.0. Although Martini-3.0 is found to be more fusogenic than Martini-2.2 (by ∼ 30 kJ mol^*−*1^),^32^ the overall effects of the non-bilayer lipids and the LHCII on the thylakoid membranes remain the same for both force fields. Thus the non-bilayer lipids assist in the formation of the non-lamellar phase mediated by fusion by reducing the Δ*G*_*stalk*_. Notably, in the presence of the LHCII, the Δ*G*_*stalk*_ of the plant thylakoid increases by 92.29 and 75.98 kJ mol^*−*1^ using the Martini-2.2 and 3.0 respectively. However, the effect of LHCII on the Δ*G*_*stalk*_ decreases with increasing distance between the protein and the stalk (data not shown). The higher Δ*G*_*stalk*_ of the plant thylakoid in the presence of LHCII is consistent with the higher *κ* obtained from the splay modulus (Figure 4 C-D inset). The Δ*G*_*stalk*_ of the plant thylakoid in the presence of the LHCII is 41 − 63 *k*_*B*_*T* which is significantly higher than the activation energy (∼ 30 *k*_*B*_*T*) of the spontaneous fusion of typical lipid vesicles.^33^ Although the nonbilayer lipids in the thylakoid assist in lamellar to non-lamellar phase transformation, LHCII provides resistance in the non-bilayer induced phase transformation by increasing the *κ* and the Δ*G*_*stalk*_ via the non-bilayer lipid redistribution from the LHCII deprived fused regions to the LHCII periphery as seen in Figure 4 E-H. The redistribution of non-bilayer lipids observed from our simulations agrees well with the earlier findings on MGDG fingerprints of LHCII trimers^34^ due to chlorophyll a^28^ and PSII^35^ as well as LHCII aggregation supported by MGDG.^36^

**Figure 5:**
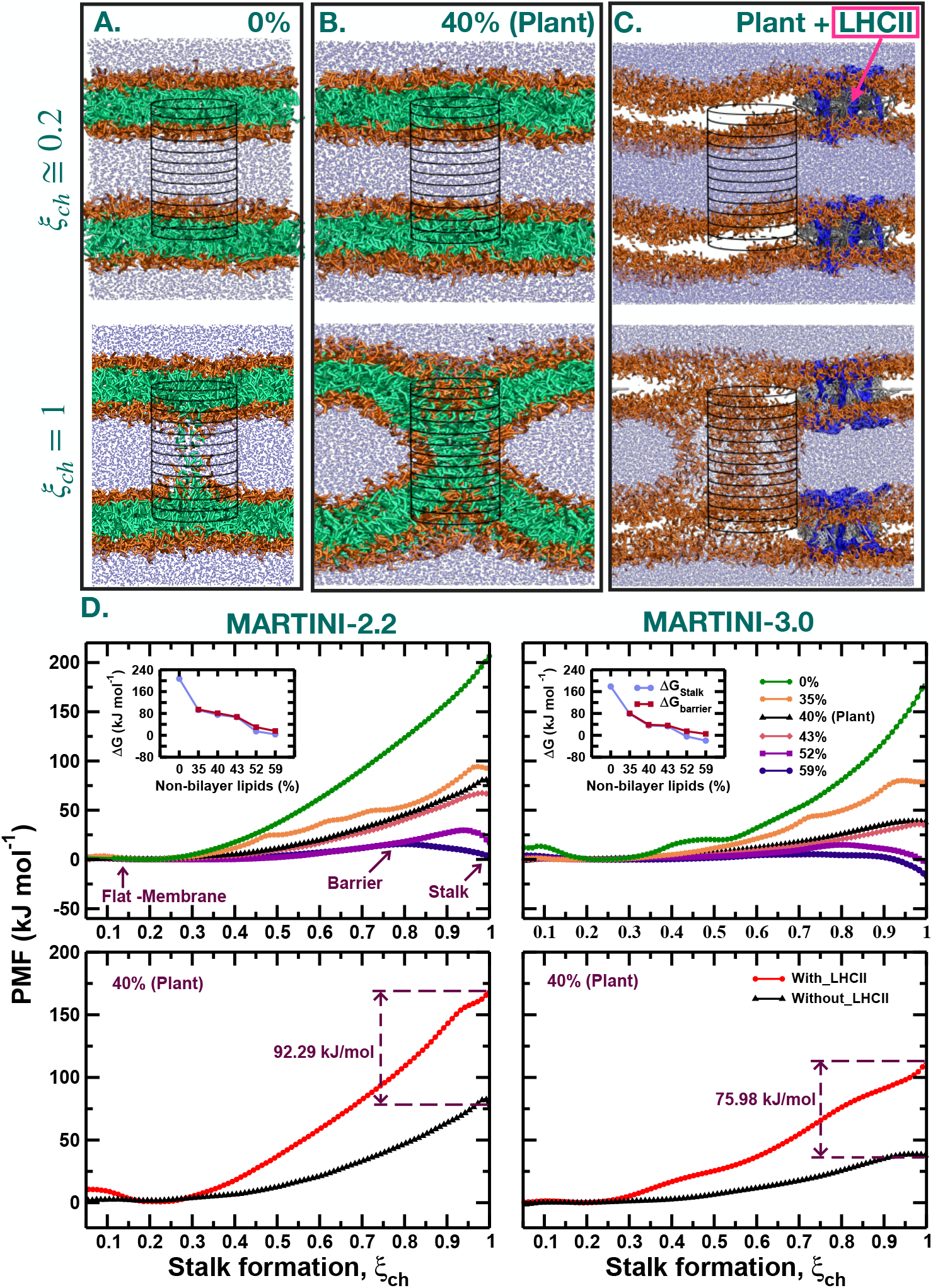
Sanpshots of double membranes with non-bilayer lipid concentrations of (A) 0%, (B) 40% without the LHCII, and (C) 40% with LHCII. The reaction coordinate, *ξ*_*ch*_, measures the connection between the two hydrophobic membrane cores during stalk formation using the membrane connecting cylinder with *N*_*s*_ slices. As the slices along the *ξ*_*ch*_ (vertically) are filled up by the lipid beads one by one, gradually a stalk is formed between two membranes. Initially, at the flat lamellar phase, a 20% of the total slices of the cylinder is filled up by lipids from both sides of the water compartment estimating *ξ*_*ch*_ = 0.2 (upper panel). When 100% slices are filled at *ξ*_*ch*_ = 1, the stalk is formed (lower panel). Color code: lipid head: orange; tails: green; water: ice-blue; LHCII trimer: blue; pigments: grey. The lipids tails are removed in (C) to show the LHCII clearly. (D) PMFs of the stalk formation of the membranes with 0 − 59% non-bilayer lipids without the LHCII (upper panel) and with 40% non-bilayer lipids and the LHCII (lower panel) using Martini-2.2 and Inset of the upper panel of D: Free energy of the stalk (Δ*G*_*stalk*_) and the barrier (Δ*G*_*barrier*_, if present) versus concentrations of the non-bilayer lipids in the thylakoid membrane, as obtained from the PMFs.

To summarize, a total of 617.8 *μ*s long unbiased and biased CG simulations using Martini-2.2 and 3.0 exhibit lipid polymorphism of the plant thylakoid membrane with a hydration level of 14.2 at 323 K in the absence of the LHCII which the LHCII resists. Undulations of the lamellar leaflets in the absence of LHCII cause interleaflet water depletion resulting in fusion that leads to a lamellar to non-lamellar phase transformation evident from the higher probability of non-zero mean curvature. Although a similar spontaneous lamellar to inverted hexagonal or cubic phase transformation of the thylakoid is found earlier at a lower temperature, it was triggered by lower hydration. ^8^ The polymorphism disappears and the lamellar phase prevails in the absence of the non-bilayer lipids. Non-bilayer lipids significantly reduce the bilayer’s bending modulus, enhancing the leaflets’ flexibility. Additionally, the free energy of stalk formation, Δ*G*_*stalk*_, decreases with increasing non-bilayer lipids, promoting fusion and nonlamellar phase transformation. Remarkably, LHCII sets off the non-bilayer lipid redistribution towards its periphery, enhancing the *κ* and the Δ*G*_*stalk*_ of the plant thylakoid and the probability of zero mean curvature for the lamellar phase becomes higher. Although Martini-3.0 is more fusogenic,^32^ the effects of the nonbilayer lipids and the LHCII on the plant thylakoid remain similar in both Martini-2.2 and 3.0. Although the thylakoids with and without the LHCII do not exhibit any phase transition under semi-isotropic pressure coupling in MARTINI-2.2, increase in *κ* and Δ*G*_*stalk*_ due to the presence of the LHCII, corroborates with the higher resistance of the LHCII to fusion for phase transition. As per our knowledge, our results, for the first time, systematically demonstrate at the molecular level that the phase transformation is assisted by the non-bilayer lipids and resisted by the LHCII leading to a trade-off between the non-bilayer lipids and the LHCII. Therefore, thylakoids can exhibit polymorphism, even with LHCII present, if the LHCII to non-bilayer lipid ratio falls below a critical threshold. These non-bilayer phases serve as an attractive site for the activity of the VDE dimer enzyme,^21^ which is essential for the xanthophyll cycle during photoprotection under excessive illumination. The extent of curvature generation in the thylakoid will depend on the assembly of the LHCII and other photosynthetic machinery. This agrees with the concept of protein-dependent nanoscopic curvature generation for cellular trafficking in eukaryotes^37^ and explains the earlier apparently contradicting observations of LHCII-driven lamellar phase of MGDG^13^ and the polymorphic state of the functional thylakoid.^21^ On a larger scale, the crowded LHCII along with its photosynthetic machinery should be capable of controlling the lipid polymorphism in thylakoid which in turn affects different biophysical processes such as NPQ through curvature-mediated activation of the xanthophyll cycle.

## Acknowledgments

A.D. acknowledges the financial support of the grant SERB CRG/2019/000106. A.G. is thankful to Tanay Sahu for the initial stage system setups.

## Supplementary information

Simulation details of unbiased runs without and with LHCII, impact of GROMACS parameters on bilayer stability, methodology to calculate mean curvature, bending modulus using height fluctuations and splay contributions, biased simulations for stalk formation free energy, and density profiles of non-bilayer lipids for the thylakoid membranes without and with LHCII.

## Notes

The authors declare no competing interest.

