## Supplementry Information for "Light Harvesting Complex II Resists Non-Bilayer Lipid-Induced Polymorphism in Plant Thylakoid Membranes via Lipid Redistribution": JPCL_Plant_LHCII_SI.pdf

### 1 Simulation Details and Methods

#### 1.1 Simulations of unbiased runs

In this study, CG MD simulations are performed on a plant thylakoid membrane consisting of seven types of lipids: $16:1(3t)-16:0$ PG, $16:1(3t)-18:3(9,12,15)$ PT, $18:3(9,12,15)-16:0$ DGDG, di$18:3(9,12,15)$ DGDT, $18:3(9,12,15)-16:0$ MGDG, di$18:3(9,12,15)$ MGDT, and $18:3(9,12,15)-16:0$ SQDG in the presence of CG water beads maintaining hydration level 14.2. The AA to CG mapping of all thylakoid lipids, LHCII cofactors chlorophyll-a, chlorophyll-b, and lutein are shown in Figure 1 of the main manuscript. Six single bilayers (S1-S6 in Table S1) are prepared using packmol[1] tool with $0-59\%$ concentrations of non-bilayer lipids, MGDG and MGDT. The bilayer with 40% non-bilayer lipids is the composition of the original plant thylakoid and is referred to as a plant throughout the manuscript. All CG simulations of S1-S6 are performed with Martini-2.2[2] and Martini-3.0[3] at a hydration level of 14.2.

Table S1: Details of bilayers for a total of 230 $\mu$s unbiased simulations using Martini-2.2 (115 $\mu$s) and Martini-3.0 (115 $\mu$s). S1-S6 are single membranes without LHCII simulated with semi-isotropic pressure coupling. F1-F6 are four stacked membranes without LHCII simulated with anisotropic pressure coupling. S3 and F3 have the compositions of the plant thylakoid. S7 and F7 are single and four-stacked plant thylakoid membranes respectively with LHCII.

| System | Number of lipids | Percentage of non-bilayer lipids | Box Size, $nm^3$ | | | NPT runlength ($\mu$s) |
|---|---|---|---|---|---|---|
| | | | x | y | z | |
| S1 | 1152 | 0 | 20.10 | 20.10 | 8.05 | 10 |
| S2 | 1824 | 35 | 24.22 | 24.22 | 8.28 | 15 |
| S3 | 1920 | 40 (Plant) | 25.08 | 25.08 | 8.44 | 10 |
| S4 | 1984 | 43 | 25.51 | 25.51 | 8.66 | 10 |
| S5 | 2208 | 52 | 26.47 | 26.47 | 8.78 | 10 |
| S6 | 2400 | 59 | 27.46 | 27.46 | 8.45 | 10 |
| S7 | 1422 | 40 (Plant+LHCII) | 22.09 | 22.09 | 9.17 | 10 |
| F1 | 1152 | 0 | 4.47 | 22.67 | 35.99 | 5 |
| F2 | 1824 | 35 | 4.39 | 29.94 | 42.96 | 5 |
| F3 | 1920 | 40 (Plant) | 4.43 | 21.96 | 57.10 | 5 |
| F4 | 1984 | 43 | 4.45 | 27.54 | 45.93 | 5 |
| F5 | 2208 | 52 | 9.03 | 12.99 | 50.51 | 5 |
| F6 | 2400 | 59 | 4.93 | 34.45 | 37.57 | 5 |
| F7 | 1656 | 40 (Plant+LHCII) | 13.52 | 11.99 | 35.59 | 10 |

Energy minimizations of the initial structures of S1-S6 are done with the steepest-descent algorithm. After that, short NVT runs are carried out for 100 ps with a time step of 10 fs at 323 K, followed by NPT simulations for each bilayer. The NPT run lengths of each system are mentioned in Table S1. The pressure is maintained at 1 bar using semi-isotropic pressure coupling by the Berendsen barostat[4] with a coupling constant of 3 ps. Coulomb interactions are calculated by the Cutoff method with a cutoff of 1.2 nm, and van der Waals interactions are treated using the cutoff scheme with a cutoff of 1.2 nm. All bonds are constrained using the LINCS algorithm.[5] Periodic boundary conditions (PBC) are applied in all three (XYZ) directions. The area per lipid of the bilayer is used to assess the equilibration of the bilayers.

The area per lipid drastically drops only for S6 using Martini-2.2 (Figure S1) and for S2-S6 using Martini-3.0, demonstrating a phase transformation. No drastic drop in area per lipid is found for S1 using Martini 2.2 and Martini-3.0. To reconfirm the phase transformation for S2-S5 obtained from Martini 3.0, stacks of four single bilayers are simulated using Martini 2.2 and Martini 3.0 which are referred to as F1-F6 in Table S1. Energy minimization of the initial structure of four stacked bilayers is done for each case with the steepest-descent algorithm. Short NVT runs are carried out for 100 ps with a time step of 10 fs. Next, each stacked bilayer is simulated at 323 K for 5 $\mu$s in the NPT ensemble using a 10 fs time step. The pressure is maintained at 1 bar using anisotropic pressure coupling by the Berendsen barostat with a coupling constant of 3 ps. The area per lipids of the stacked bilayers F2-F6 in Martini-2.2 reduces with $1 - 2$ $\mu$s, indicating similar phase transformations of F2-F6 as observed using Martini-3.0 (Figure S1). The reduction in area per lipid is more drastic in Martini-3.0 than in MARTINI-2.2, suggesting that the phase transformation is comparatively favorable in MARTINI-3.0.

### 1.2 Simulations of unbiased runs in the presence of LHCII

To study the effect of the LHCII on the polymorphism of the plant thylakoid membranes, LHCII is simulated in the presence of a thylakoid bilayer with 40% non-bilayer lipids. The initial all-atom (AA) configuration of the LHCII having the protein trimer, chlorophyll-a, chlorophyll-b, and lutein molecules is obtained from the protein data bank (1RWT.pdb).[8] The AA protein trimer is mapped into a CG protein configuration using the program martinize.[9] The AA co-factors of LHCII are mapped into CG representation using VOTCA[10] following mapping schemes of both Martini-2.2[2] and Martini-3.0.[3] The ElNeDyn[11] model for only Martini-2.2 and a standard elastic network model for Martini-3.0 were used to maintain the secondary and tertiary protein structure. The energy-minimized pigment binding protein trimer CG LHCII is embedded in a plant thylakoid bilayer using the Python tool insane.[12]

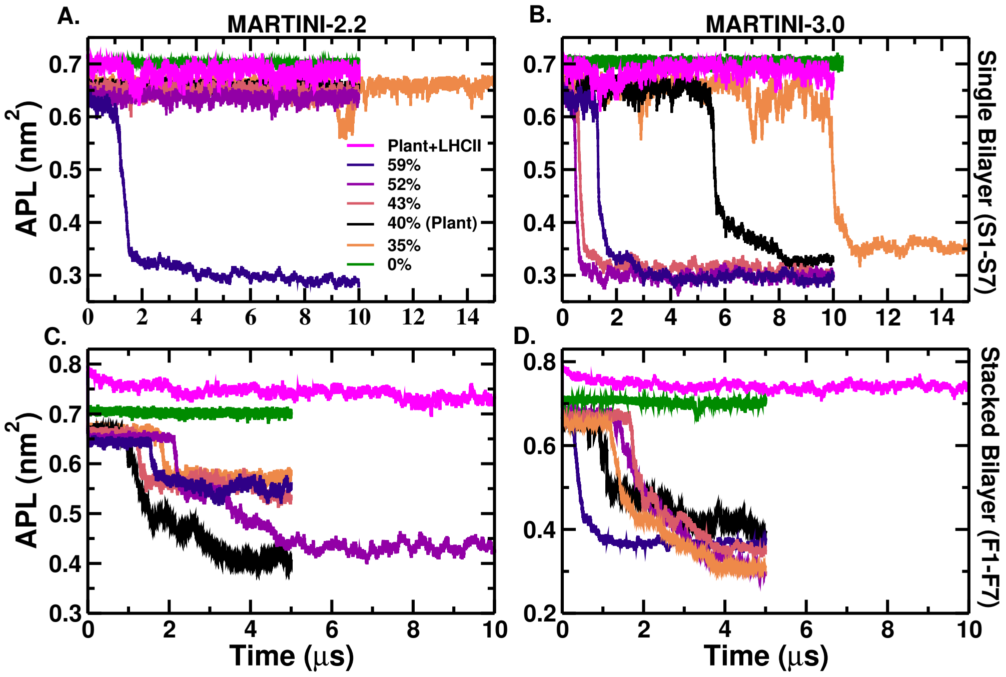


Figure S1: Area per lipid of (a) S1-S7 in Martini-2.2 with semi-isotropic pressure coupling (b) S1-S7 in Martini-3.0 with semi-isotropic pressure coupling (c) F1-F7 in Martini-2.2 with anisotropic pressure coupling. (d) F1-F7 in Martini-3.0 with anisotropic pressure coupling. The $a_h$ of the plant thylakoid without the LHCII at the lamellar state from our simulation is $\sim 0.65 \pm 0.0065$ nm$^2$ for Martini-2.2 and 3.0 at 323 K which agrees reasonably well with the previously reported $a_h$ $0.66 \pm 0.003$ $nm^2$ of the bilayer at 293 K.[6]

The LHCII embedded bilayer is solvated with 20027 CG water beads, and 562 positively charged sodium ions are added to neutralize the protein. An energy minimization is carried out, followed by a short 100 ps NVT runs with a time step of 10 fs. For all backbone beads of LHCII trimer protein, a position restraint of 4000 kJ mol$^{-1}$ nm$^{-2}$ is applied for a 10 ns NPT run, then position restraint is reduced to half for another 10 ns NPT run, and finally, a position restraint of 200 kJ mol$^{-1}$ nm$^{-2}$ is used for the 10 $\mu$s NPT run. After that position restarined is removed for the 10 $\mu$s NPT run with semi-isotropic pressure coupling. The LHCII embedded plant thylakoid is referred to as S7 in Table S1 where the system details are mentioned. The four stacked bilayers system with the LHCII, referred to as F7 in Table S1, is simulated for a 10 $\mu$s NPT run with anisotropic pressure coupling using both Martini-2.2[2] and Martini-3.0.[3] All other simulation parameters are the same as the ones used in the above simulation details. All unbiased simulations are performed using GROMACS version

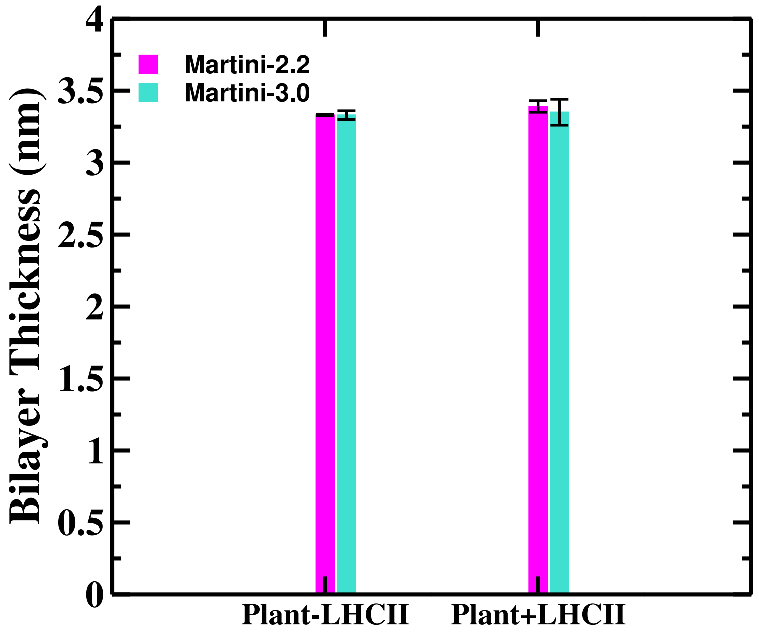


Figure S2: Bilayer thickness of the plant thylakoid membrane is $3.33 \pm 0.005 - 0.03$ nm and $3.35 - 3.39 \pm 0.09 - 0.04$ nm without and with the LHCII respectively using Martini-2.2 and 3.0. This is similar to the thickness of the thylakoid with and without PsbS as $\sim 4.3$ and 3.2 nm respectively upon illumination.[7]

2018.1.

### 1.3 Impact of GROMACS Parameters on Bilayer Stability

To understand the effect of GROMACS parameters on our results, four stacked plant thylakoid bilayers have been simulated for 500 ns with Martini-3.0 using semi-isotropic pressure couping with two conditions: the default nstlist = 10 with Verlet-buffer-tolerance default (VBT = 0.005 kJ mol$^{-1}$ ps$^{-1}$) and nstlist = 1 with Verlet-buffer-tolerance (VBT = $-1$ kJ mol$^{-1}$ ps$^{-1}$). Deviations of the diagonal pressure tensor elements from the target pressure in MD simulations with the nstlist = 1 with default (VBT = 0.005 kJ mol$^{-1}$ ps$^{-1}$) and nstlist = 10 with VBT = $-1$ kJ mol$^{-1}$ ps$^{-1}$ are shown in Figure S3. The area per lipid (APL) of the lamellar phase is similar for both simulations, however, they are different for the non-lamellar phase for both simulations (as seen in Figure S4). The drop in area per lipid occurs at similar time scales for both cases. This suggests that the lamellar to non-lamellar phase transformations in both cases occur in a similar time scale; the geometries of non-lamellar phases are minorly affected by the choice of nstlist and VBT which is reflected in Figure S5. However, the deviations of the pressure tensors from the reference pressure do not suppress the undulation in our case for both simulations. This is an indicator that the geometries of the non-lamellar phase might slightly vary on the choices of nstlist and VBT, the observed undulations in our simulations are physical and independent of GROMACS parameters and not an artifact.

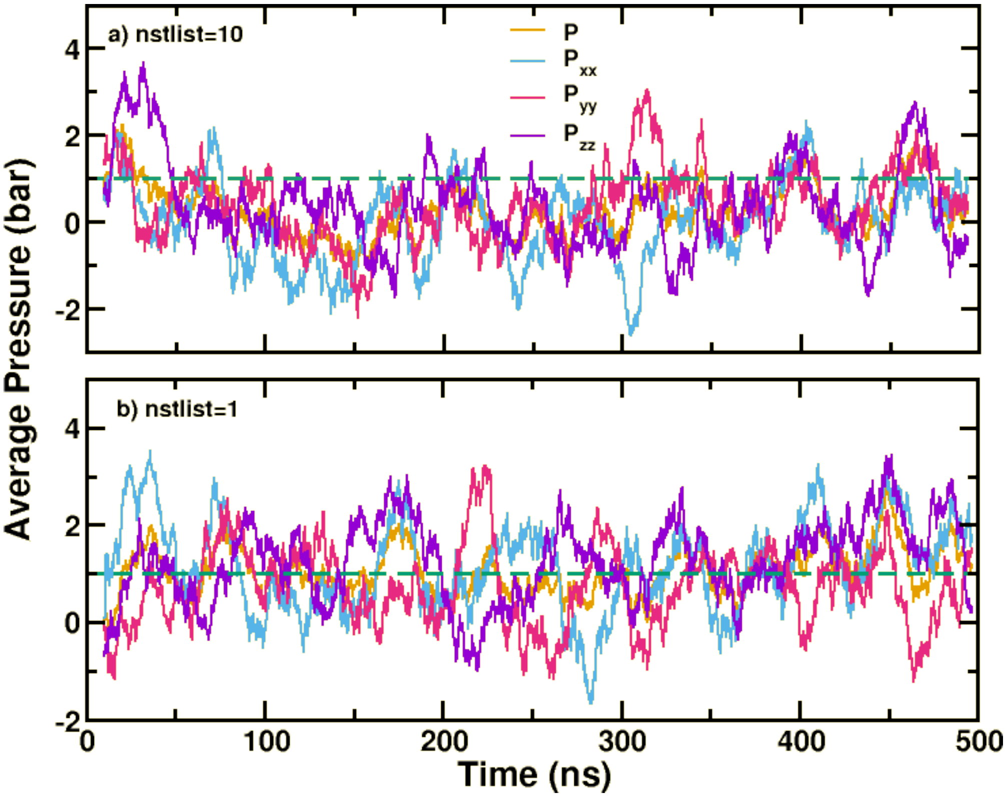


Figure S3: Running average of diagonal pressure tensor elements and the target pressure in MD simulations with the (a) nstlist=10 with default (VBT = 0.005 ) and (b) nstlist=1 with VBT=−1 kJ $mol^{-1}$ $ps^{-1}$.

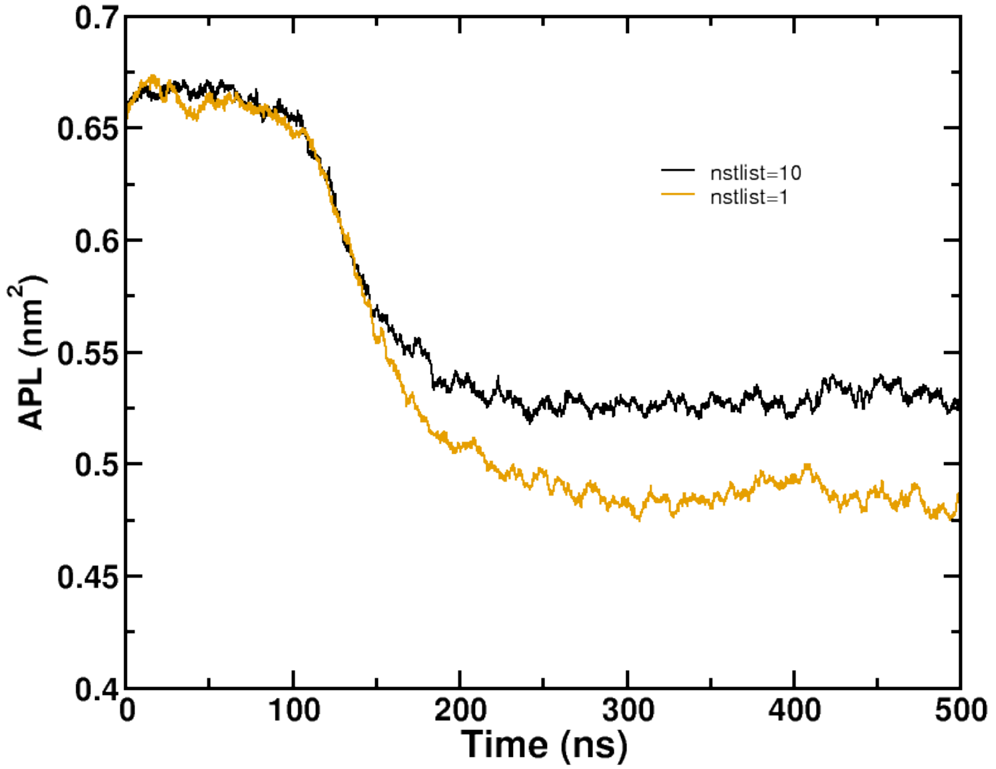


Figure S4: Area per lipid of large stacked bilayer system using the nstlist=10 with default (VBT = 0.005) and nstlist=1 with VBT=−1 kJ mol$^{-1}$ ps$^{-1}$.

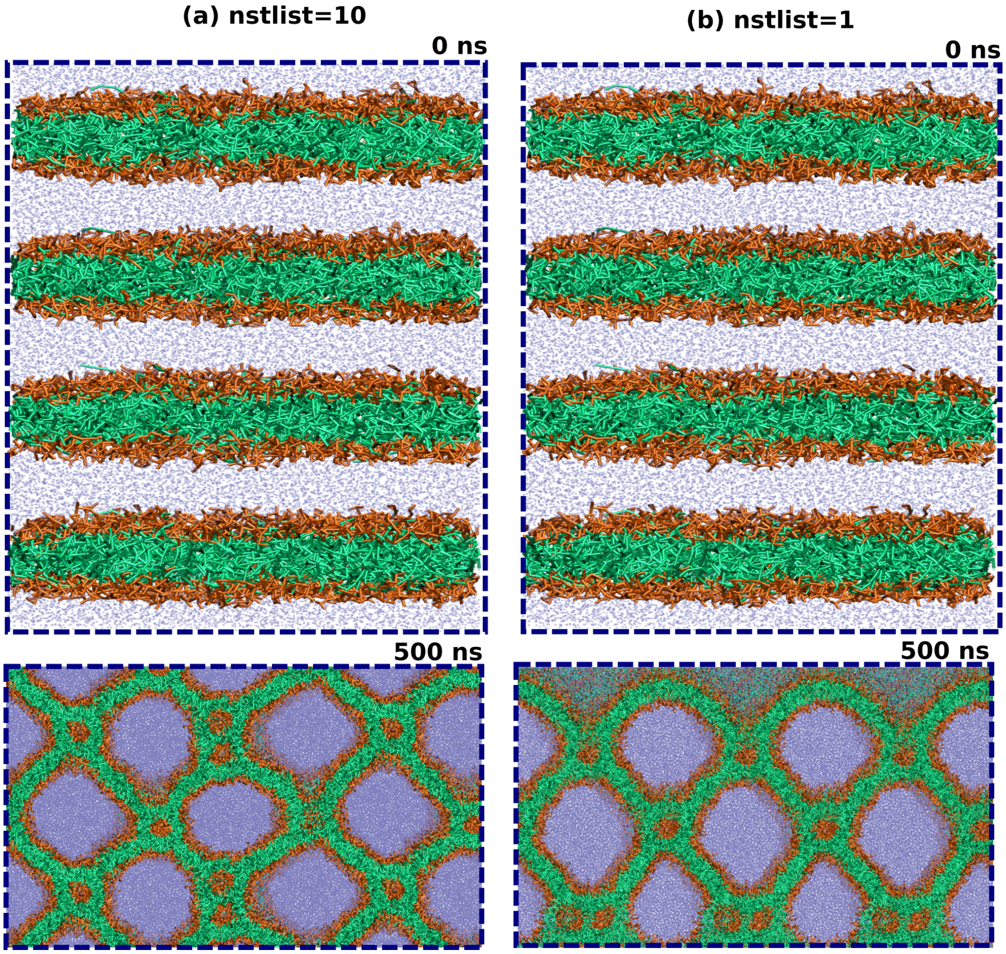


Figure S5: Snapshot of a stacked bilayer ($25 \times 25$ nm$^2$ in x-y) at 0 ns and after fusion at 500 ns with (a) nstlist = 10 with default (VBT = 0.005 kJ mol$^{-1}$ ps$^{-1}$) and (b) nstlist = 1 ( VBT=$-1$ kJ mol$^{-1}$ ps$^{-1}$).

### 1.4 Mean Curvature ($H$)

To understand the probability of non-lamellar surfaces formed for the plant thylakoid membranes in the presence and absence of the LHCII, 10 and 20 configurations of the plant thylakoid membranes are generated using the random velocity generation method in the presence and absence of LHCII respectively (see Table S2) using MARTINI-3.0 and semi-isotropic pressure coupling. Similar analyses are not performed with MARTINI-2.2 since no phase transition is observed. Each thylakoid membrane is simulated for a 10 $\mu s$ NPT run. The entire 10 $\mu s$ for each replica are utilized to calculate the probability distributions of the mean curvature using the method mentioned in.[13] The custom software for calculating membrane mean curvature is available from (https://github.com/bio-phys/MemCurv).

Table S2: Details of a total of 300 $\mu$s (out of this 20 $\mu$s is already reported in Table S1) unbiased simulations using Martini-3.0 of different configurations of the plant thylakoid bilayer in the presence and absence of LHCII for calculating $H$.

| System | Number of lipids | Number of replicas | NPT run-lengths ($\mu$s) of each replica |
|---|---|---|---|
| Plant-LHCII (S3) | 1824 | 20 | 10 |
| Plant+LHCII (S7) | 1152 | 10 | 10 |

### 1.5 Bending Modulus Calculation ($\kappa$)

Further, the locally equilibrated run-lengths before the phase transformation of S1-S6 are used to calculate the bending modulus of the lamellar phase from the height fluctuations[14] and from the splay contributions.[15] Additionally, the last 1 $\mu$s run-length after the phase transformation is chosen to calculate the bending modulus of the fused state from splay contributions. The last 1 $\mu$s run-length of 10 $\mu$s run is used to calculate the bending modulus of S7.

#### 1.5.1 Using height fluctuation

The first method to calculate the bending modulus is using a Fourier transformation of the height fluctuations $h(x, y)$ of the membrane surface. 10000 frames are used to calculate the $h(x, y)$ for each of the lamellar phase of S1-S6. The Fourier transform of $h(x, y)$ results in $h(q)$ where $q$ is a wave vector. The intensity spectrum, $S(q)$, of the height fluctuations can be represented by the following equations using the Helfrich model of the bilayer.

$$S(q) =< |h(q)|^2 A) > \quad (1)$$

The angular bracket represents the time average, and $A$ is the surface area. To determine the bending modulus, one measures the power spectrum of the height fluctuations over a wide range of $q$. The log-log plot of $S(q)$ versus $q$ in the low $q$ regime is expected to follow $q^{-4}$ scaling, and the bending modulus ($\kappa$) is determined from the intercept (see equation 2 in the main manuscript) of the plot. At small $\gamma$ in the regime of small $q$, the undulatory mode dominates over the protrusion mode for a tension-less bilayer. However, to obtain reliable results, a sufficiently large system size must be used to ensure the system is tension-free. This is because the tension in the membrane can affect the measurements and skew the results. Figure S3 illustrates the $S(q)$, as a function of $q$, for systems S1-S6 using both force fields Martini-2.2 and Martini-3.0. The figure shows that $S(q)$ decreases as the $q$ increases, which is a common trend in systems with a single bilayer. The solid line in the figure corresponds to the $q^{-4}$ fit at low frequencies. The depicted $q^{-4}$ fitting parameters of Figure S6 are mentioned in Tabel S3.

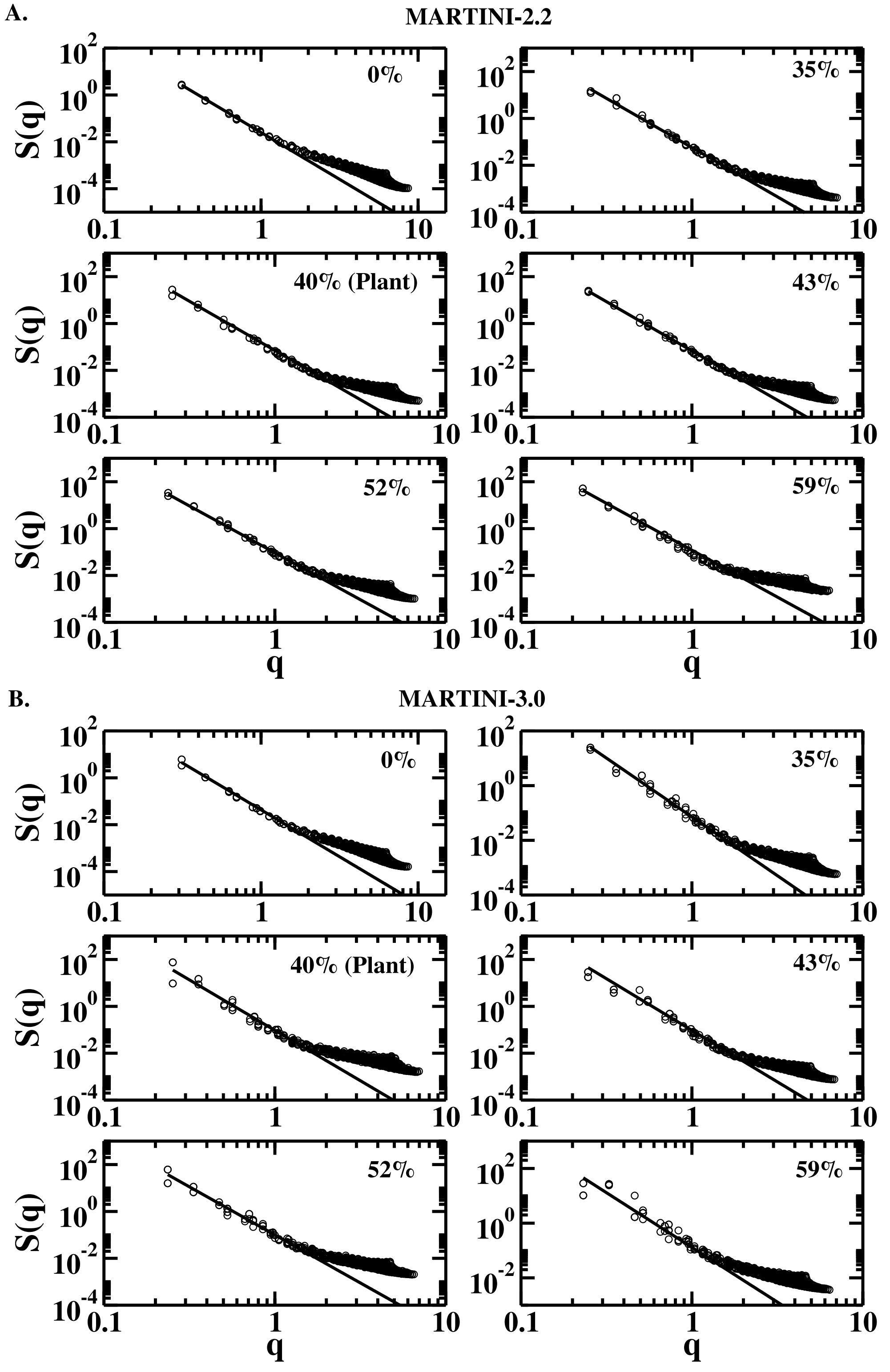


Figure S6: Intensity spectrum of S1-S6. The solid line corresponds to a $q^{-4}$ fit and illustrates that $q^{-4}$ scaling is captured in all cases. Furthermore, the $q^{-4}$ scaling observed in the low $q$ range indicates that the bending modes have a greater influence than the protrusion modes.

Table S3: Fitting parameters of $S(q)$ vs $q^{-4}$ in $k_BT$.

| System | Percentage of non-bilayer lipids | $1/\kappa$ from fitting ($k_BT$). MARTINI-2.2 | MARTINI-3.0 |
|---|---|---|---|
| S1 | 0 | 0.025 | 0.043 |
| S2 | 35 | 0.057 | 0.072 |
| S3 | 40 (Plant) | 0.071 | 0.089 |
| S4 | 43 | 0.088 | 0.095 |
| S5 | 52 | 0.903 | 0.102 |
| S6 | 59 | 0.119 | 0.130 |

#### 1.5.2 Using splay modulus ($\chi_{12}$) contribution

The splay modulus $\chi_{12}$ is calculated by quadratic fitting of $PMF(\alpha)$:

$$\begin{aligned} PMF(\alpha) &= \chi_{12}\ \alpha^2 + c \\ &= -k_BT\ ln\frac{P(\alpha)}{sin\alpha} \end{aligned} \tag{2}$$

Combined probability distribution $P(\alpha)$ of all pairs of thylakoid lipids (of S1-S7) are plotted as a function of the splay angle $\alpha$ between the lipids. The splay angle ($\alpha$) is calculated for only those pairs of lipids where at least one of them is tilted at an angle of no more than $\theta = 10^o$ concerning the bilayer normal (Figure S4). To narrow down the study to nearby molecules, only pairs of lipids within a distance of 2 nm from each other are included in these computations. The 5000 frames from each run of the S1-S7 are used for this calculation. The data presented are calculated using a coarse-grained approach for the S1-S7 systems in the lamellar phase (Figure S5) and S2-S6 systems for the fused phase (Figure S6). The custom software for calculating membrane bending modulus from splay contribution is available from (https://github.com/cpc1996/Kc-Bending-Modulus/tree/main).

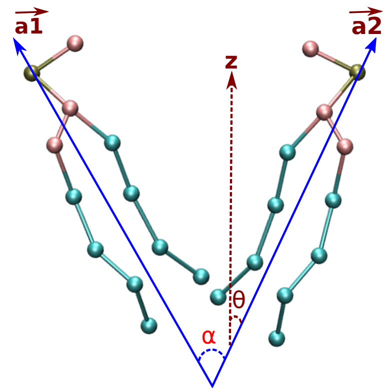


Figure S7: Schematic representation of lipid-lipid pair (in bonded representation) showing the definitions used for tilt and splay angle calculations. The vectors $a1$ and $a2$ along two lipids make the splay angle $\alpha$. $\theta$ denotes the respective tilt angles with respect to the bilayer normal z.

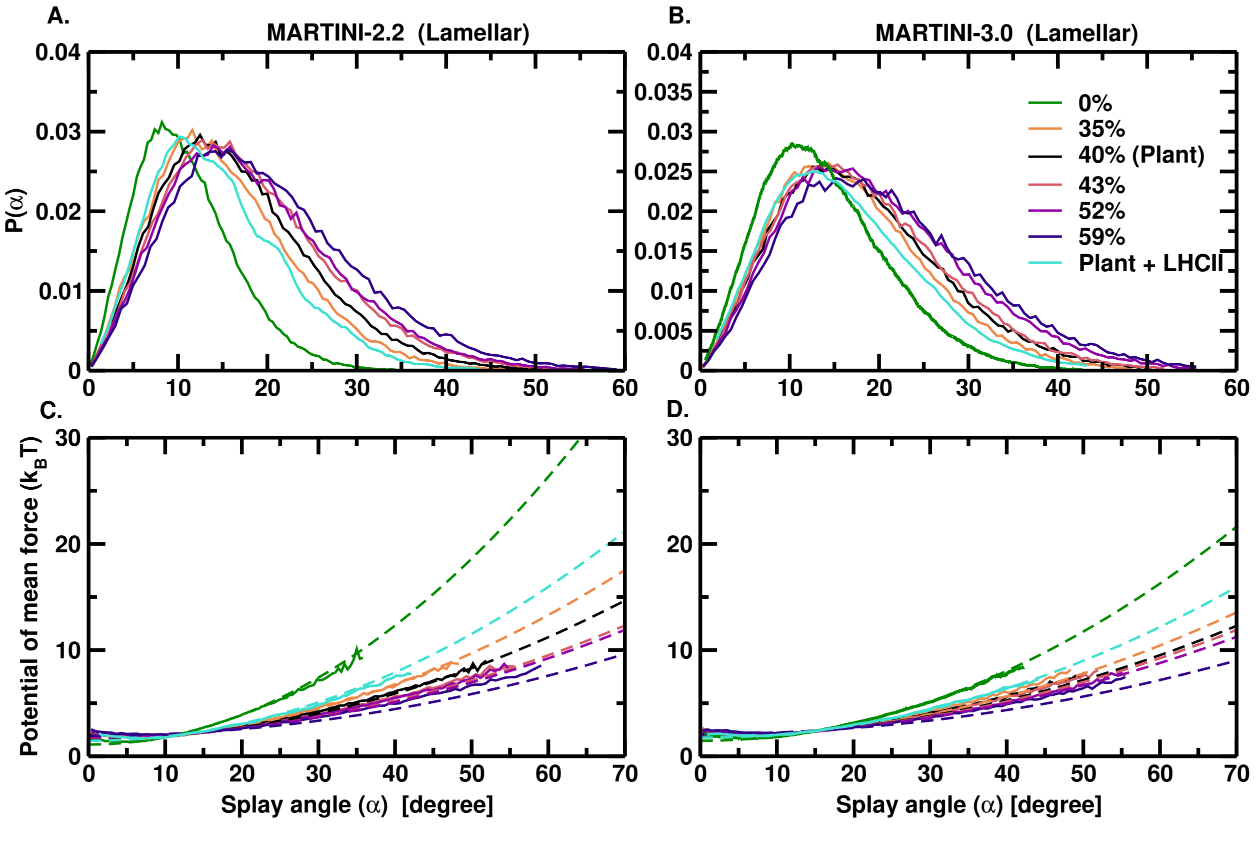


Figure S8: (Top) Combined normalized probability distribution $P(\alpha)$ of all pairs of thylakoid lipids for each system (in S1-S7) in lamellar phase. (Bottom) The potential of mean force profiles ($PMF(\alpha)$, shown by solid curves. The dashed lines show the most accurate quadratic fittings, from which the associated splay modulus $\chi_{12}$ are computed. Quadratic fitting is applied within the range of $\alpha$ values from $5-20$ for the lamellar phase. This range is chosen to focus on the low-angle regime while still incorporating the most well-sampled regions in the $P(\alpha)$ distribution profiles.

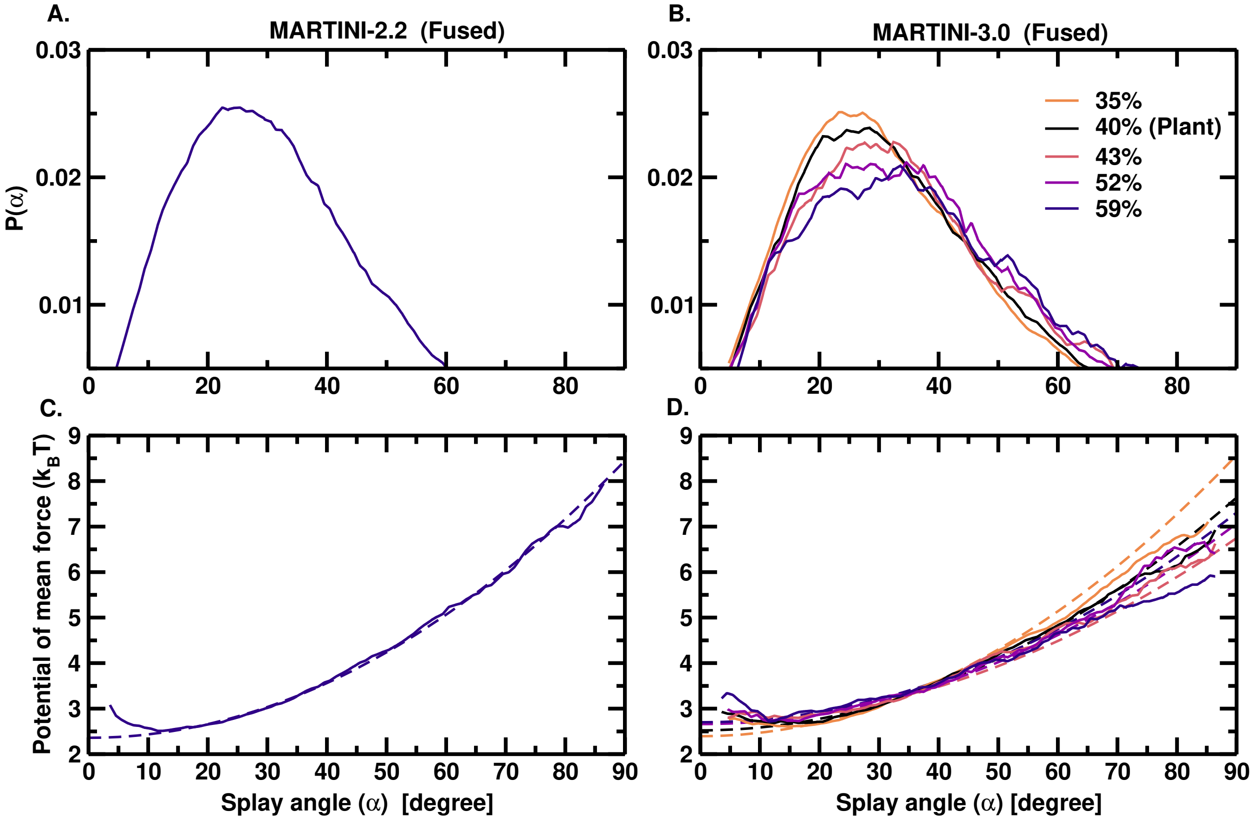


Figure S9: (Top) Combined normalized probability distribution $P(\alpha)$ of all pairs of thylakoid lipids for each system (in S2-S6) in the fused phase. (Bottom) The potential of mean force ($PMF(\alpha)$ profiles, shown by solid curves. The dashed lines show the most accurate quadratic fittings, from which the associated splay modulus $\chi_{12}$ are computed. Quadratic fitting is applied within the range of $\alpha$ values from $10 - 40$ for the fused phase.

Table S4: The bending modulus, $\kappa$ of the pre-equilibrated[18] AA DMPC bilayer from our calculations using height fluctuations and splay contributions align well with the reported value from computation[16] and experiment,[17] validating our methods.

| System | Bending modulus, $\kappa$ ($k_BT$) | |
|---|---|---|
| DMPC | Height | Splay |
| (303K) | Fluctuation | Contribution |
| AA (Our simulation) | 29.88 | $30.05 \pm 2.09$ |
| AA (Literature)[16] | | 34.71 |
| Experiment[17] | 13.29-31.10 | |

Table S5: Bending Modulus ($\kappa$) obtained from the unbiased MD runs of S1-S7 using both the methods of height fluctuations and splay contributions for the lamellar phase. For the non-lamellar fused state, splay contributions are used to calculate the $\kappa$. No position restraint is applied with the protein.

| System | Percentage of non-bilayer lipids | Bending modulus, $\kappa$ ($k_BT$) | | | | | |
|---|---|---|---|---|---|---|---|
| | | MARTINI-2.2 | | | MARTINI-3.0 | | |
| | | Height Fluctuation (Lamellar) | Splay Contribution (Lamellar) | Splay Contribution (Fused) | Height Fluctuation (Lamellar) | Splay Contribution (Lamellar) | Splay Contribution (Fused) |
| S1 | 0 | 39.97 | $41.25 \pm 0.31$ | - | 23.25 | 23.53±0.29 | No fusion |
| S2 | 35 | 17.33 | 17.89±0.63 | - | 13.88 | 13.63±0.65 | 3.74±0.25 |
| S3 | 40 (Plant) | 14.08 | 14.30±0.69 | - | 11.23 | 11.50±0.65 | 3.46±0.32 |
| S4 | 43 | 11.27 | 11.64±0.77 | - | 10.52 | 10.88±0.71 | 3.19±0.31 |
| S5 | 52 | 11.01 | 10.84±0.76 | - | 9.76 | 9.86±0.43 | 3.18±0.33 |
| S6 | 59 | 8.39 | 8.33 ±0.77 | 4.38±0.18 | 7.69 | 7.24±0.76 | 2.84±0.34 |
| S7 | 40 (Plant+LHCII) | - | $21.65 \pm 0.49$ | - | - | $14.96 \pm 0.76$ | - |

### 1.6 Simulation of stalk formation and umbrella sampling

To calculate the free energy of stalk formation from umbrella sampling simulations,[19] a biased potential is applied along the chain reaction coordinate, $\xi_{ch}$[20] generally used to observe pore formation in the membrane.[21] The biased harmonic potential applied is as follows:

$$\omega_i(\xi_{ch}) = \frac{k}{2}(\xi_{ch} - (\xi_{ch}^{ref})_i)^2 \tag{3}$$

Here $\omega_i$ represent biasing potential, $(\xi_{ch}^{ref})_i$ is the reference point for respective window $i$. $k$ is the strength of the bias, and $\xi_{ch}$ is the chain reaction coordinate.

The potential of mean force[22] is calculated by

$$PMF(\xi_{ch}) = -\int_{-\infty}^{\infty} \left\langle \left( \frac{\partial V}{\partial q_{\xi_{ch}}} \right)_{q\{m \not\equiv \xi_{ch}\}^{N-1}} \right\rangle_{\xi_0} dq^{N-1}_{\{m \not\equiv \xi_{ch}\}} \tag{4}$$

where, $q$ represents the configuration, $V$ is the potential energy, and $N$ is the dimension of the phase space.

For defining the $\xi_{ch}$, a cylinder of radius 1.2 nm spans the regions of double membranes and the middle water compartment between two thylakoid membranes, and the cylinder is made up of $N_s$ slices of thickness 0.1 nm. $\xi_{ch}$ is given by the fraction of slices filled with lipids beads $n_s(t)$ defined as below:

$$\xi_{ch} = \frac{1}{N_s} \sum_{N_s}^{N_s-1} \delta_{\xi}(n_s(t)) \tag{5}$$

$N_s$ is the number of cylinder slices, and $n_s(t)$ is the number of lipid beads in slice s. The differential indicator function, $\delta_{\xi}$, $(0 < \delta_{\xi} \leq 1)$, assumes a value 0 if no beads are present in slice $s$, and takes a value close to 1 if slices are filled. The formulated $\delta_{\xi}$ is defined with a differentiable switch functions as follows:

$$\delta_{\xi}(x) = \begin{cases} \zeta_x & if \quad x \leq 1 \\ 1 - ce^{-kx} & if \quad x > 1 \end{cases} \tag{6}$$

The parameter $\zeta$ denotes the fraction to which a slice is filled upon adding the lipid bead into the slice. A typical value for $\zeta$ would be 0.75. The parameters $b$ and $c$ are taken as $b = \zeta/(1-\zeta)$ and $c = (1-\zeta)e^b$, leading to the continuous and differentiable switch function. Likewise, the number of lipid beads $n_s(t)$ located in slice $s$ is formulated in terms of switch functions as follows:

$$n_s(t) = \sum_{i=1}^{N^{(b)}} f(r_i, z_s, d_s, X_{cyl}, Y_{cyl}, R_{cyl}) \tag{7}$$

Here, $r_i = (x_i, y_i, z_i)$ are the cartesian coordinates of atom $i$, and $N^{(b)}$ is the number of lipids beads in the entire system. The indicator function $f$ takes unity if bead $i$ is located within slice $s$ and a cylinder of radius $R_{cyl}$, and it takes zero otherwise. The lateral position of the cylinder in the membrane plane is generally dynamically defined, to allow the cylinder to follow the stalk. However, a dynamic cylinder in the presence of the protein allows a stalk formation to happen away from the protein which does not give the correct estimate of the $\Delta G_{stalk}$ due to the protein. Thus, the cylinder is kept fixed near the protein to get the maximum impact of the protein to the $\Delta G_{stalk}$. This results in the formation of a double stalk (Figure 5C of the main text) near the protein since the protein does not facilitate the formation of the stalk. During pulling the system along $\xi_{ch}$, the slices are filled with lipid beads one by one, gradually forming a stalk between two membranes. Here, the $\xi_{ch} \approx 0.2$ corresponds to a flat unperturbed membrane, implying that 20% of the cylinder slices are filled by lipid beads from both sides of the water compartment. $\xi_{ch} \approx 1$ corresponds to a fully formed stalk.

Two membranes are stacked along the z-direction from the energy-minimized configuration of a single bilayer with $0-59\%$ of non-bilayer lipids. These systems are referred to as D1-D6 in Table S6. The stacked bilayers with the plant thylakoid composition in the absence and presence of LHCII are referred to as D3 and D7 respectively.

Table S6: Details of bilayers and a total of 107.8 $\mu s$ biased simulation runs for PMF calculations. D1-D6 are two stacked membranes containing variable concentrations of non-bilayer lipids without LHCII and D7 is the plant thylakoid with LHCII.

| System | Number of lipids | Percentage of non-bilayer lipids | Stalk formation (chain reaction coordinate) Pulling (ns) | Equilibration runlength(ns) | Production runlength(ns) | Number of windows |
|---|---|---|---|---|---|---|
| D1 | 576 | 0 | 200 | 200 | 200 | 19 |
| D2 | 912 | 35 | 200 | 200 | 200 | 19 |
| D3 | 960 | 40 (Plant) | 200 | 200 | 200 | 19 |
| D4 | 992 | 43 | 200 | 200 | 200 | 19 |
| D5 | 1104 | 52 | 200 | 200 | 200 | 19 |
| D6 | 1200 | 59 | 200 | 200 | 200 | 19 |
| D7 | 828 | 40 (Plant+LHCII) | 200 | 200 | 200 | 19 |

The double membrane for each case is energy minimized using the steepest descent algorithm. Next, 100 ps NVT runs are carried out, followed by a 50 ns NPT run to relax the box dimensions fully. The final configuration of the 50 ns NPT runs are the starting configurations for pulling simulation using chain reaction coordinates.[23] The initial geometries for umbrella sampling simulations are taken from the pulling simulation, in which the system is pulled from $\xi_{ch} \approx 0.2$ to $\xi_{ch} = 1$ within 200 ns. The pulling force constant of 5000 kJ mol$^{-1}$ nm$^{-2}$ and a pull rate of $4.5 \times 10^{-6}$ nm ps$^{-1}$ are used for constant-velocity pulling simulations. For PMF calculation, 19 umbrella windows are used from the reference position of 0.1 to 1 in steps of 0.05. Each window is simulated for 400 ns, where the first 200 ns are omitted for equilibration. The weighted histogram analysis method (WHAM)[24] is employed to calculate the PMF from the biased simulation windows. All other parameters are kept the same as the ones used in the unbiased simulation details. When the cylinder for the stalk formation is kept dynamic near the LHCII, the stalk is formed away from the protein which results in much reduced $\Delta G_{stalk}$. In the absence of LHCII, the non-bilayer lipids are located in the fused region whereas in the presence of the LHCII, these lipids are in the annular region of the trimer providing rigidity near the protein and resistance to fuse. As one goes away from the LHCII, this effect weakens and $\Delta G_{stalk}$ decreases. Although there is a reduction in the $\Delta G_{stalk}$, the addition of the LHCII increases the value of $\Delta G_{stalk}$ by $4 - 19$ $k_BT$ for MARTINI-3.0 and MARTINI-2.2 respectively, even when the cylinder is dynamic. This again suggests that the LHCII resists stalk formation in the membrane. The overlapping of histograms in Figure S7 shows a good sampling over the entire chain reaction coordinate. The PMFs (Figure S8) of stalk formation of plant thylakoid with LHCII (D7) using Martini-2.2 either in stalk-opening direction or in stalk-closing direction are similar. This is a signature of no hysteresis in chain reaction coordinates. The error bars suggest that the PMFs are converged. To apply GROMACS Chain Coordinate, patched GROMACS version 2021.6 is utilized for all biased runs.

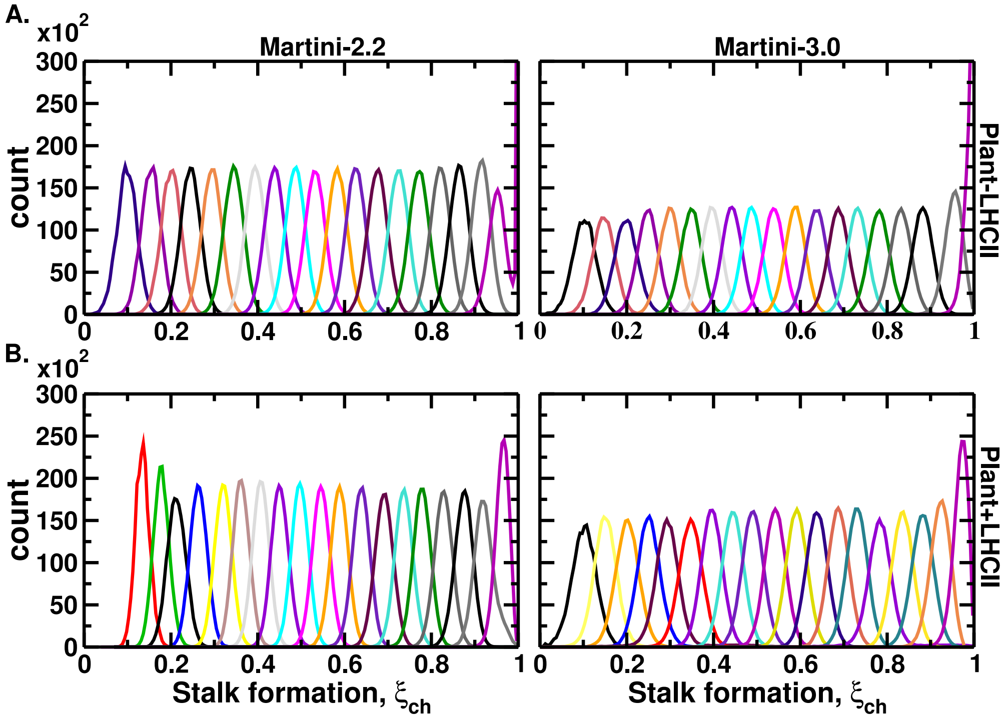


Figure S10: Histograms of 40% plant thylakoid membrane in the (a) absence (D3) and (b) presence (D7) of LHCII using Martini-2.2 and using Martini-3.0.

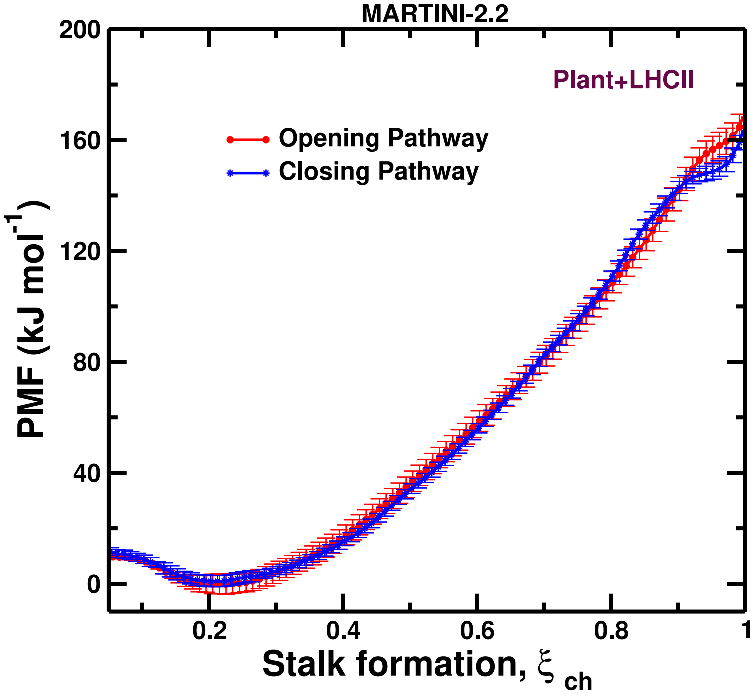


Figure S11: PMFs of stalk formation of plant thylakoid with LHCII (D7) using Martini-2.2 either in stalk-opening direction (blue) or in stalk-closing direction (red). Similar PMFs for the opening and closing suggest the absence of any hysteresis and error bars suggest that the PMFs are converged.

Table S7: Free energies $\Delta G_{stalk}$ of stalk formation with varying concentrations of non-bilayer lipids using umbrella sampling simulations. The $\Delta G_{stalk}$ for the plant thylakoid without LHCII is 74.97 kJ $mol^{-1}$ for Martini-2.2 at 323 K, aligning well with the inner leaflet of the plasma membrane with a similar density of water beads per $nm^2$.[20]

| System | Percentage of non-bilayer lipids | MARTINI-2.2 $\Delta G_{stalk}$ (kJ/mol) | MARTINI-2.2 $\Delta G_{barrier}$ (kJ/mol) | MARTINI-3.0 $\Delta G_{stalk}$ (kJ/mol) | MARTINI-3.0 $\Delta G_{barrier}$ (kJ/mol) |
|---|---|---|---|---|---|
| D1 | 0 | 207.26 | - | 178.77 | - |
| D2 | 35 | 92.57 | 94.40 | 78.79 | 80.24 |
| D3 (323K) | 40 (Plant) | 74.97 | 81.09 | 35.96 | 38.93 |
| Plasma membrane[20] (inner leaflet) (310 K) | | ≈ 75 | | | |
| D4 | 43 | 66.59 | 67.42 | 33.07 | 35.93 |
| D5 | 52 | 14.58 | 29.68 | -4.34 | 14.82 |
| D6 | 59 | 3.33 | 15.64 | -19.35 | 5.29 |
| D7 | 40 (Plant+LHCII) | 168.83 | - | 111.94 | - |
